# Maternal Effects and Parent-Offspring Conflict

**DOI:** 10.1101/168310

**Authors:** Bram Kuijper, Rufus A. Johnstone

**Affiliations:** Environment and Sustainability Institute (ESI), College of Life and Environmental Sciences, University of Exeter, Penryn Campus, Penryn TR10 9EZ, UK; Department of Zoology, University of Cambridge, Downing Street, Cambridge CB2 3EJ, UK

**Keywords:** Nongenetic effects, maternal hormone, transgenerational effect, inheritance, information, epigenetics

## Abstract

Maternal effects can provide offspring with reliable information about the environment they are likely to experience, but also offer scope for maternal manipulation of young when interests diverge between parents and offspring. To predict the impact and outcome of parent-offspring conflict, we model the evolution of maternal effects on local adaptation of young. We find that parent-offspring conflict strongly influences the stability of maternal effects; moreover, the nature of the disagreement between parents and young predicts how conflict is resolved: when mothers favour less extreme mixtures of phenotypes relative to offspring (i.e., when mothers stand to gain by hedging their bets), mothers win the conflict by providing offspring with only limited amounts of information. When offspring favour overproduction of one and the same phenotype across all environments compared to mothers (e.g., when offspring favour a larger body size), neither side wins the conflict and signaling breaks down. Only when offspring favour less extreme mixtures relative to their mothers (the case we consider least likely), offspring win the conflict and obtain full information about the state of the environment. We conclude that a partial or complete breakdown of informative maternal effects will be the norm rather than the exception in the presence of parent-offspring conflict.

## 1 Introduction

Maternal effects comprise any causal influence of the environment or phenotype of the mother on the phenotype of her offspring that is not mediated by genetic transmission (Wolf & Wade, 2009; Day & Bonduriansky, 2011; Danchin *et al.*, 2011). Such effects have been identified in many species, and may involve a wide variety of different mechanisms, ranging from hormonal influences (von Engelhardt & Groothuis, 2011), through the transmission of antibodies (e.g., Boulinier & Staszewski, 2008) and maternal provisioning of nutrients (e.g., Wells, 2010), to social learning (Mesoudi *et al.*, 2016) and even active teaching (Rapaport, 2011). It is well established that maternal effects can, at least in principle, strongly influence the course of evolution within a population (Mousseau & Fox, 1998; Räsänen & Kruuk, 2007; Badyaev & Uller, 2009; Hoyle & Ezard, 2012). More recently, there has been much discussion of when and why selection might favour the evolution of such effects themselves (Kuijper *et al.*, 2014; English *et al.*, 2015; Kuijper & Hoyle, 2015; McNamara *et al.*, 2016; Proulx & Teotόnio, 2017).

Adaptive explanations of the evolution of maternal effects often suggest that they serve to provide offspring with information about the environment they are likely to encounter (Marshall & Uller, 2007; Shea *et al.*, 2011; Kuijper & Johnstone, 2013; Leimar & McNamara, 2015). This information allows the young to anticipate the challenges they will face and to develop an appropriate phenotypic response (Agrawal *et al.*, 1999; Galloway & Etterson, 2007; McGhee & Bell, 2014; Holeski *et al.*, 2012, but see Uller *et al.*, 2013). For instance, offspring field crickets (*Gryllus pennsylvanicus*) born from mothers that have been exposed to predators exhibit greater antipredator immobility (Storm & Lima, 2010). Other antipredator adaptations have been observed in *Daphnia*, where offspring from mothers that have been exposed to predatory stimuli grow larger defensive helmets (Agrawal *et al.*, 1999). Similar processes also operate in plants, for example in *Campanulastrum americanum*, where offspring phenotypes are dependent on the maternal light environment, and those that experience a light environment that matches that of their mother have a 3.4 times larger fitness in comparison to offspring that develop in different light conditions (Galloway & Etterson, 2007). These examples show that in at least some cases, maternal effects facilitate offspring anticipation of environmental challenges.

Maternal effects, however, do not always operate to the advantage of offspring. In some cases, they appear to benefit the mother at the expense of individual young (Jaenike, 1986; Einum & Fleming, 2000; Mayhew, 2001; Janz *et al.*, 2005; McCormick, 2006, reviewed in Marshall & Uller, 2007). For example, in *Cephaloleia* beetles, maternal survival is increased when ovipositing on novel plant hosts, whereas individual offspring survival was reduced compared to young on native hosts (García-Robledo & Horvitz, 2012). This raises the question whether mothers always stand to gain by supplying information beneficial to their young.

Parents and offspring often face an evolutionary conflict of interest (Trivers, 1974; Parker & Macnair, 1978; Godfray, 1995; Smiseth *et al.*, 2008; Kilner & Hinde, 2008). This conflict arises because offspring value their own survival more strongly than that of current or potential future siblings, while parents value all of their offspring equally (Trivers, 1974). Behavioural ecologists have focused mostly on conflicts over resource provisioning, in which offspring are selected to demand more resources than parents are selected to provide (Parker & Macnair, 1978; Godfray, 1995; Hinde *et al.*, 2010; Wells, 2007a,b; Kuijper & Johnstone, 2012). However, this conflict may influence information exchange as well. In particular, much attention has been devoted to parents’ acquisition of information about offspring need or hunger, and to what extent they can rely on offspring signals of condition (Godfray, 1991; Godfray & Johnstone, 2000; Royle *et al.*, 2002; Wells, 2003). Here, by contrast, we are concerned with acquisition of information about the environment by offspring from their parents, but similar issues arise within each context of information exchange (Uller & Pen, 2011). When there is parent-offspring conflict over the optimal offspring phenotype, can offspring rely on maternal signals about the environment? Alternatively, might maternal effects provide a means by which mothers can manipulate offspring phenotype and enforce their own optima on their young (Müller *et al.*, 2007; Uller, 2008; Kilner & Hinde, 2008; Tobler & Smith, 2010)?

So far, how parent-offspring conflict affects the evolution of informative maternal effects has seen surprisingly little formal investigation. A single model by Uller & Pen (2011) has considered how parent-offspring conflict over dispersal affects the degree of information contained in maternal effects. Unless offspring are somehow constrained in their response to maternal signals, they find that parent-offspring conflict typically does not affect the evolution of informative maternal effects, so that at evolutionary equilibrium, offspring are able to rely on maternal signals to implement their own optimal strategy. This contrasts markedly with other signalling models that focus on information transfer from offspring to parents, which suggest that conflict leads to the breakdown of informative signalling, unless honesty is maintained by some form of signal cost (Godfray *et al.*, 1991; Johnstone, 1999; Godfray & Johnstone, 2000). Consequently, this raises the question of whether informative signalling by mothers to offspring is indeed a general outcome of parent offspring conflict, as suggested by Uller & Pen (2011), or whether there are contexts in which conflict can lead to a breakdown of informative maternal signals instead.

To assess how parent-offspring conflict affects the evolution of maternal effects, we focus on a scenario of conflict over offspring local adaptation in a spatiotemporally varying environment (Leimar & McNamara, 2015; English *et al.*, 2015; Kuijper & Johnstone, 2016). Fluctuating environments often favour parents that produce a mixture of offspring phenotypes, containing some offspring that are adapted and some offspring that are maladapted to the current state of the local environment (Starrfelt & Kokko, 2012). Producing a mixture of offspring phenotypes ensures that at least some offspring are likely to survive, even if the local environment changes, thus preventing the extinction of the parental gene lineage (Ellner, 1986; McNamara, 1995; Leimar, 2005). In contrast to their parents, however, individual offspring have a higher genetic interest in their own survival than in that of their siblings. Consequently, offspring favour a lower probability of developing a currently maladapted phenotype than do their parents, resulting in parent-offspring conflict over local adaptation (Ellner, 1986).

We explore a situation in which offspring cannot assess the environment they will experience directly for themselves, but must rely on signals from their mother. A key ingredient of our model is that mothers can potentially ‘skew’ the information they provide, by signalling in a misleading way. The question we then seek to answer is whether reliable maternal signalling is stable, allowing for the persistence of maternal effects, or whether it is vulnerable to disruption by maternal dishonesty.

## 2 The model

We consider an ‘infinite island’ model (Wright, 1931; Rousset, 2004; Lehmann & Rousset, 2010) comprising a sexually hermaphroditic metapopulation that is distributed over an infinite number of patches, each of which contains *n* adult breeders. Generations are discrete and non-overlapping, and in each generation, every breeder produces, as mother, a large number of offspring, each of which is sired by a random breeder. With probability *ℓ*, this sire is chosen from the same patch as the mother (including the possibility of self-fertilisation), while with probability 1 – *ℓ* the sire is chosen from a random remote patch. For the sake of tractability, we assume that the population is haploid, where gametes are produced clonally and pair to form diploid zygotes, which immediately undergo meiosis to form a new generation of haploid offspring (individual-based simulations assuming diploid inheritance and a finite number of patches give similar results, see Figures S6-S8). Upon birth, a fraction 1 – *d* of newborn young remain on the natal patch, while the remaining fraction *d* disperse to a random patch in the metapopulation. After dispersal, offspring on a patch, both native and immigrant, compete for the *n* breeding vacancies created by the death of the previous generation. Those that fail to obtain a breeding vacancy die, and the life cycle then repeats. Below we provide a verbal summary of the model, while a more extensive description is given in section S2 of the Online Supplement.

**Environmental variation** Following previous models of maternal influences on offspring phenotype determination that do not consider parent-offspring conflict (e.g., Shea *et al.* 2011; English *et al.* 2015; Leimar & McNamara 2015; Kuijper & Johnstone 2016), we consider a spatiotemporally fluctuating environment in which each patch fluctuates between two environmental states, *e*_1_ and *e*_2_. In each generation, an *e_i_* patch can change to an *e_j_* patch with probability *σ*_*i*→*j*_ (*i* ≠ *j*) while it remains in environmental state *e_i_* with probability 1 – *σ*_*i*→*j*_. Patches fluctuate independently of one another, so that at any given time a proportion *p*_1_ = *σ*_2→1_/(*σ*_1_→_2_ + *σ*_2→1_) of patches is in environmental state *e*_1_, while the remainder *p*_2_ ≡ 1 – *p*_1_ is in environmental state *e*_2_.

**Phenotype determination** Upon birth of an offspring, it can adopt one of two phenotypes, *z*_1_ or *z*_2_. Individuals are ‘locally adapted’ and therefore experience a lower mortality rate when their phenotype *z_i_* is identical to the environment *e_i_* of their patch (Kawecki & Ebert, 2004). Individuals are characterised by the genetically determined strategy *f_i_* which reflects the probability that an offspring develops phenotype *z*_1_ as opposed to phenotype *z*_2_. Importantly, *f_i_* may depend upon an offspring’s natal environment *e_i_*, so we consider the evolution of a strategy **f** = {*f*_1_, *f*_2_} that specifies phenotype determination probabilities for each of the two environments. Our model also accounts for the possibility that offspring of one phenotype are potentially more costly to the mother (i.e. they require more maternal resources) than offspring of the opposite phenotype (e.g., Trivers, 1974; Ellner, 1986; Kuijper & Pen, 2014). Moreover, we allow such maternal production costs to vary dependent on the local environment *e_i_*, so that the parameters *β_i_* and *γ_i_* reflect the maternal cost of producing a *z*_1_ and *z*_2_ offspring respectively when the local environment is in state *e_i_.* Hence, the average investment *E_i_* by a mother living in environment *e_i_* per offspring is proportional to *f_i_β_i_* + (1 – *f_i_*) *γ_i_.* Following classical life-history models (Smith & Fretwell, 1974; Parker & Macnair, 1978), we assume that the total number of offspring produced is inversely proportional to the average investment per offspring. Consequently, the proportions of *z*_1_ and *z*_2_ offspring produced by a mother living in environment *e_i_* are then given by *f_i_*/*E_i_* and (1 – *f_i_*)/*E_i_* respectively. After phenotype determination, offspring either disperse or stay in the local patch, with dispersal occurring prior to environmental change. The survival probability of an offspring with phenotype *z_j_* that ends up competing in a patch that is in environmental state *e_j_* is given by *ω_ij_.* Throughout, we assume that offspring with a phenotype that matches the local environment always survive, so that *ω*_11_ = *ω*_22_ = 1, while *z*_1_ offspring in an *e*_2_ environment survive with probability *ω*_12_ =1 – *c*_2_ and *z*_2_ offspring in an *e*_1_ environment survive with probability *ω*_21_ = 1 – *c*_1_. All surviving offspring in a patch, both immigrant and philopatric, then compete for the *n* adult breeding positions that are locally available. The resulting fitness equations, which describe the number of successfully established offspring born from adults living in each environment are set out in section S2.1 of the Online Supplement.

### 2.1 Mapping the battleground

The question now arises to what extent the evolutionary interests of parents and offspring diverge when it comes to the decision of developing phenotype *z*_1_ versus *z*_2_. To resolve this issue, we compare the evolutionarily stable values of *f*_1_ and *f*_2_ under maternal and under offspring control (the divergence between these outcomes defining the ‘battleground’ within which parent-offspring conflict will be played out, Godfray, 1995). To determine the equilibrium probabilities of producing a *z*_1_ phenotype under either maternal or offspring control, we adopt an adaptive dynamics approach (Geritz *et al.*, 1998; Rousset, 2004; Dercole & Rinaldi, 2008). This assumes that evolution proceeds by the successive substitution of mutations of small effect, with a clear separation of time scales between demographic and evolutionary processes (Otto & Day, 2007). We use a direct fitness (also called neighbour-modulated fitness) approach (Taylor & Frank, 1996; Taylor *et al.*, 2007) to derive the selection gradient ℱ_*i*_ that determines the evolutionary change in the probability *f_i_* of producing a *z*_1_ offspring in environment *e_i_* (see eq. [S5]). By numerically iterating the selection gradients until they vanish, we are able to solve numerically for the equilibrium probabilities
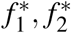
of producing a *z*_1_ phenotype in each of the two environments.

### 2.2 Resolving the conflict

If the interests of mothers and offspring diverge, how then might maternal-offspring conflict be resolved? If offspring must rely on mothers for information about the state of the local environment, could this enable mothers to manipulate the behaviour of their young in each of the two contexts considered? To evaluate this possibility, we suppose that mothers can assess the state of the local environment, while offspring cannot (see also Uller & Pen, 2011). Mothers may choose to give or to withhold a signal from each of their young, with probabilities of giving the signal dependent on the state of the local environment. Offspring may then choose to develop phenotype *z*_1_ or *z*_2_, with probabilities dependent on whether or not they have received a signal from their mother.

The maternal signalling strategy **s** ≡ (*s*_1_, *s*_2_) thus specifies the probabilities of giving (rather than withholding) the signal in each type of patch, while the offspring phenotype determination strategy **q** = (*q_S_*, *q_NS_*) specifies the probabilities of developing a *z*_1_ phenotype when a signal is or is not received. It is the combination of these two strategies that determines the fraction of young *f_i_* that develop as *z*_1_ in each environment *e_i_*:

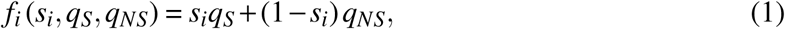

so that with probability *s_i_* a mother living in environment *e_i_* provides her offspring with a signal, who will therefore develop as a *z*_1_ offspring with probability *qs* (and as a *z*_2_ offspring with probability 1 – *qs*). By contrast, with probability 1 – *s_i_*, the mother withholds the signal, in which case offspring develops as a *z*_1_ or *z*_2_ offspring with probabilities *qNS* and 1 –*qNS* respectively. Associated fitness expressions for the maternal signaling-probabilities and offspring phenotype determination strategies are given in eqns. (S17 - S20) in the Online supplement. We again assume that evolution proceeds by the successive substitution of mutations of small effect, with a clear separation of time scales between demographic and evolutionary processes (Otto & Day, 2007). This allows us to use a direct fitness approach to derive the selection gradients 𝒮_*i*_ and 𝒬_*j*_ that determine the rates of evolutionary change in the probability *s_i_* of providing offspring a signal in each environment and the probability *q_j_* ∈ {*q_S_*, *q_NS_*} of producing a *z*_1_ offspring in the presence or absence of a signal (see eqns. [S21,S22]).

To solve the conflict resolution model, we seek to identify equilibrium strategy pairs for which all selection gradients (for both strategies) are simultaneously equal to zero. To do so, we choose initial conditions such that the signal is highly informative (e.g., we might choose *s*_1_ = 0.9 and *s_2_* = 0.1) and offspring highly responsive (e.g., *f_s_* = 0.9 and *f_NS_* = 0.1), and iteratively update the signalling and phenotype determination probabilities by adding to each the value of the relevant selection gradient (given the current strategies), bounding the updated values between 0 and 1. This procedure is repeated until all strategies converge to stable values. The solutions obtained in this way are robust to changes in the precise starting conditions chosen, and convergence stable by construction. Note, however, that two mirror-image signalling equilibria are possible in any particular case – one in which the signal is given more often in environment *e_1_* and withheld more often in environment *e*_2_, and one in which the signal is given more often in environment *e*_2_ and withheld more often in environment *e*_1_. These provide offspring with equal information, and thus have identical consequences in terms of the phenotype determination rates out of each patch type. For ease of interpretation, however, we consistently choose starting conditions in which the signal is given more often in environment *e*_1_. Individual-based simulations, which assume a continuous distribution of mutations and no necessary separation of timescales, yield very similar results to the analytical model (see Figures S6-S8).

## 3 Results

### 3.1 The battleground

Figure 1 illustrates the ways in which the interests of mothers and offspring diverge. The graphs show the stable fraction of *z*_1_ offspring produced in environment *e*_1_ (blue) and in environment *e*_2_ (red), under maternal control (dotted lines) versus under offspring control (solid lines), as a function of *c*_2_, the cost of maladaptation in environment *e*_2_ (while holding *c*_1_, the cost of maladaptation in environment *e*_1_, constant at 0.8). In general, both mothers and offspring favour higher proportions of *z*_1_ offspring when the cost of maladaptation in environment *e*_2_ is low (at the left-hand side of each graph), and lower proportions of *z*_1_ offspring when the cost of maladaptation in environment *e*_2_ is high (at the right-hand side of each graph). However, stable outcomes under maternal versus offspring control rarely agree precisely.

**Figure 1.**
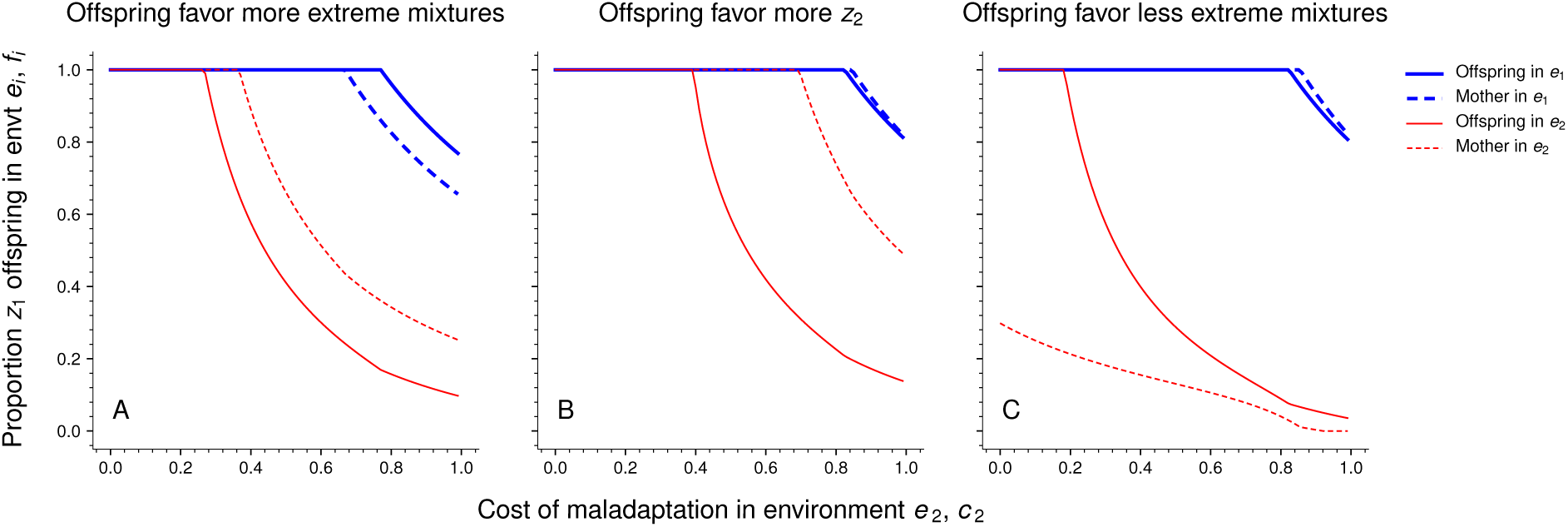
Stable probabilities of producing a phenotype *z*_1_ offspring in environments *e*_1_ (blue lines) and *e*_2_ (red lines) respectively, plotted against the cost of maladaptation *c*_2_ in environment *e*_2_. Panel A: both offspring phenotypes are equally costly to produce to mothers (*β*_1_ = *β*_2_ = *γ*_1_ = *γ*_2_ = 1). Consequently, mothers (dashed lines) favour more even mixtures of offspring phenotypes. By contrast, offspring favour more extreme mixtures that are biased towards the phenotype with the highest survival in the local environment (i.e., offspring favour more *z*_2_ in environment *e*_2_ and more *z*_1_ in environment *e*_1_). Panel B: when phenotype *z*_2_ is more costly to produce in both environments (*β*_1_ = *β*_2_ = 1, *γ*_1_ = *γ*_2_ = 2), the probability of producing *z*_2_ offspring is reduced. However, as offspring are more related to themselves than to their mothers, offspring favour a greater probability of producing more costly *z*_2_ offspring in both environments. Panel C: Maternal production costs are environment dependent, so *z*_2_ young are more costly (less costly) to produce than *z*_1_ young in environment *e*_1_ (in environment *e*_2_); (*β*_1_ = 1, *β*_2_ = 2, *γ*_1_ =2, *γ*_2_ = 1). Consequently, mothers favour more extreme mixtures of offspring phenotypes that are biased towards the phenotype that is cheaper to produce in each environment. By contrast, offspring favour a larger probability of developing as the more costly phenotype, leading them to favour more even mixtures of offspring phenotypes. Parameters: *d* = 0.1, *ℓ* = 0.5, *σ*_12_ = 0.2, *σ*_21_ = 0.25, *n* =1, *c*_1_ = 0.83.

The three panels of the figure show results for three different sets of parameter values, which we have chosen to illustrate three possible kinds of ‘disagreement’ between mother and young (see Supplementary Figure S1 for a more extensive overview of model results).

Scenario 1 (panel A): offspring favour production of more of the locally adapted phenotype in each environment (i.e. more of phenotype *z*_1_ in environment *e*_1_, and more of phenotype *z*_2_ in environment *e*_2_); in terms of the graph, the red and blue solid lines for equilibria in the case of offspring control lie ‘outside’ the corresponding dotted lines for equilibria in the case of maternal control. In this scenario, *z*_1_ and *z*_2_ offspring are equally costly to produce. Under these circumstances, mothers do best (in either environment) to hedge their bets by producing a certain fraction of young with a currently (locally) maladapted phenotype, to ensure survival of at least some of their brood in case the environment changes. Since offspring, by contrast, have a greater evolutionary interest in their own survival than in that of the brood as a whole, they favour a higher probability of developing the currently well-adapted phenotype.

Scenario 2 (panel B): Offspring favour production of more of phenotype *z*_1_ across both environments; in terms of the graph, the red and blue solid lines for equilibria in the case of offspring control lie above the corresponding dotted lines for equilibria in the case of maternal control. In this case, *z*_2_ offspring are twice as costly for mothers to produce as are *z*_1_ offspring. Under these circumstances, mothers favour mixtures of offspring phenotypes that are more biased towards the cheaper *z*_1_ phenotype across both environments, because producing a larger fraction of costly young reduces their overall fecundity. By comparison, offspring are less concerned with maternal fecundity relative to their own survival, and so favour mixtures of phenotypes that are more biased towards the expensive *z*_2_ phenotype, across both environments.

Scenario 3 (panel C): Offspring favour production of more of the locally maladapted phenotype in each environment (i.e. more of phenotype *z*_2_ in environment *e*_1_, and more of phenotype *z*_1_ in environment *e*_2_); in terms of the graph, the red and blue solid lines for equilibria in the case of offspring control lie ‘inside’ the corresponding dotted lines for equilibria in the case of maternal control. In this case, maternal costs of producing one phenotype versus the other are assumed to depend on the local environment: specifically, we assume that a *z*_2_ young is twice as costly to produce as a *z*_1_ young in environment *e*_1_, while *z*_1_ young are twice as costly to produce than *z*_2_ young in environment *e*_2_. In this case, mothers favour more extreme mixtures that are biased towards the the locally-adapted phenotype that is the cheapest to produce in that particular environment, while offspring favour less extreme mixtures that feature more of the locally costly phenotype.

We have chosen parameter values to highlight the different kinds of conflict that can arise between mothers and young, because the nature of the ‘disagreement’ turns out to affect the resolution of the conflict, as detailed below.

### 3.2 Resolution of the conflict

How is parent-offspring conflict resolved when offspring control the determination of their phenotype, but must rely on maternal signals about the state of the local environment? We can categorise outcomes of the model according to the extent of information supplied by mothers to their young - offspring may obtain (i) full information about the environment (because the presence or absence of the maternal signal is perfectly correlated with the state of the environment), (ii) partial information (because the signal is given more commonly in one environment than in the other, but the correlation is imperfect) or (iii) no information (because the presence or absence of the signal is uncorrelated with the environment).

Alternatively, taking into account both the probabilities of the signal being given or withheld, and the response of offspring in each case, we can categorise outcomes according to the degree to which the realised probabilities of producing each phenotype match the values favoured by mothers versus young - (i) parents may win (i.e. the outcome matches what evolves under maternal control), (ii) neither ‘side’ may win (i.e. the outcome diverges from what evolves under either maternal or offspring control), or (iii) offspring may win (i.e. the outcome matches what evolves under offspring control). Figure 2 shows the regions of parameter space in which the model predicts different levels of information transfer, while Figure 2 shows the regions in which mothers or offspring (or neither) are predicted to win (with equivalent results for additional regions of parameter space shown in Supplementary Figures S3 and S4).

**Figure 2.**
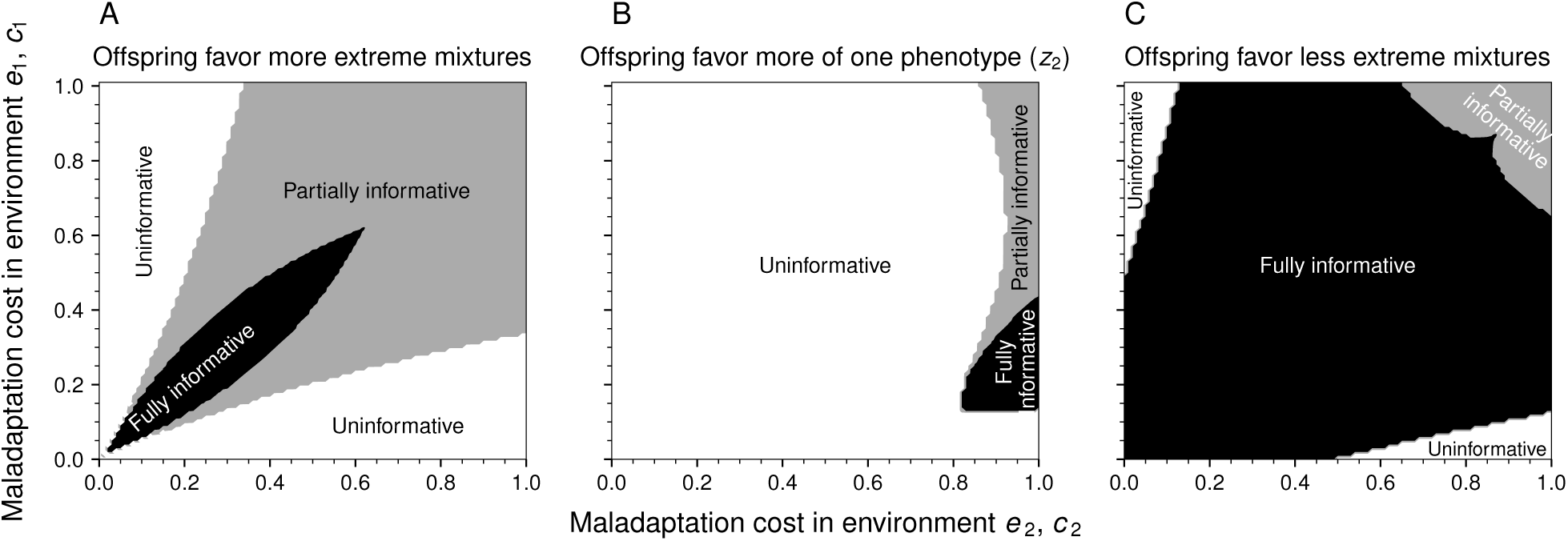
Phenotype determination when offspring rely on the maternal signal: the information content of the maternal signal **s** = (*s*_1_, *s*_2_) for the three different scenarios of conflict considered in Figure 1. Panel A: offspring phenotypes are equally costly to produce to mothers. For a wide range of costs of maladaptation, mothers evolve signals that are partially informative to offspring, although other outcomes also occur. Panel B: when the *z*_2_ phenotype is more costly to produce for mothers in both environments, maternal signals always evolve to be uninformative when conflict occurs. Panel C: when the *z*_1_ and *z*_2_ phenotypes are more costly to produce in the respective environments *e*_2_ and *e*_1_, maternal signals typically evolve to be fully informative, apart from a narrow boundary in which signals are only partially informative. The information content *H* of the signal is calculated as a measure of entropy weighed by the probability of receiving and not receiving a maternal signal in both environments:
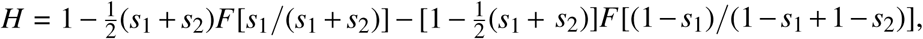 where *F*[*x*] = –*x*log_2_(*x*) – (1 – *x*)log_2_(1 – *x*). See Figure S2 for the corresponding equilibrium probabilities of producing offspring with phenotype *z*_1_ when offspring rely on a maternal signal. In addition, Figure S3 plots outcomes for an asymmetric environment where *e*_1_ patches are more common than *e*_2_ patches. Parameters: *d* = 0.1, *ℓ* = 0.5, *σ*_12_ = 0.15, *σ*_21_ = 0.15, *n* = 1.

As detailed below, comparison of Figures 2 and 3 with Figure 1 reveals that there is not necessarily a strict relationship between the nature of the parent/offspring battleground and the outcome (in terms of either information conveyed or who wins the battle). At the same time, however, there is a strong correlation, such that each of the three battleground scenarios we list above is typically associated with a different kind of outcome.

**Figure 3.**
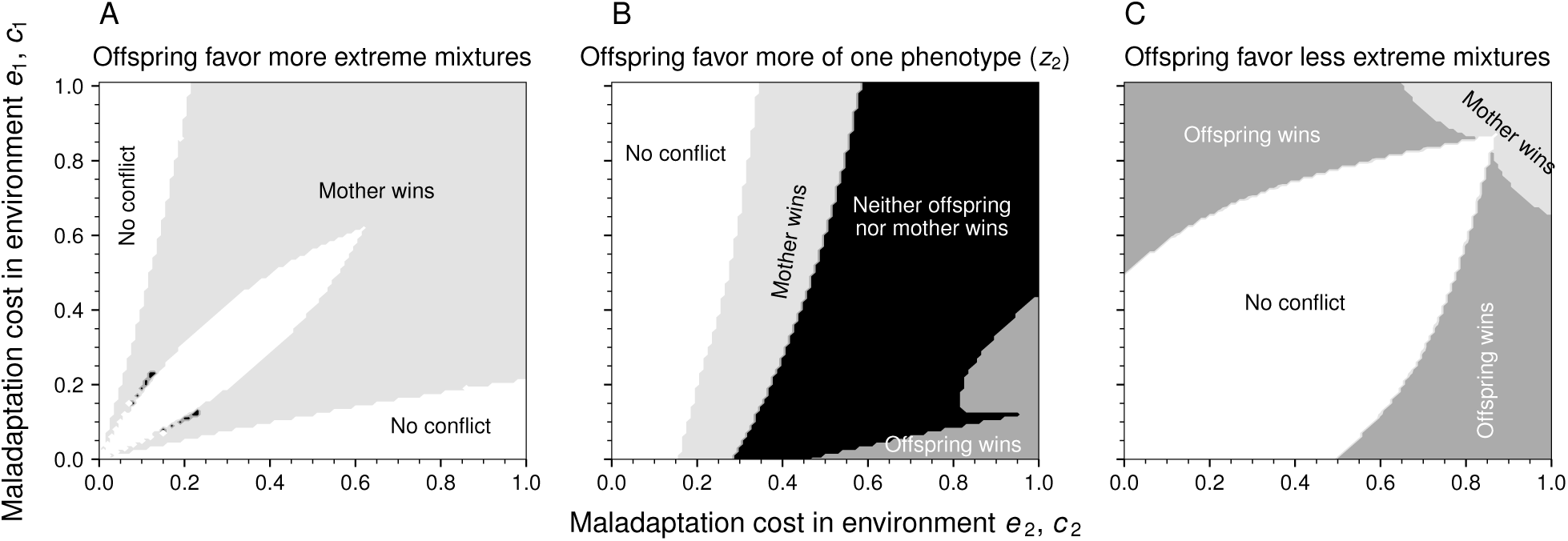
Phenotype determination when offspring rely on the maternal signal: who wins the conflict? Panel A: when offspring favour more extreme mixtures of phenotypes than mothers, mothers can be said to win the conflict by restricting the information content of the maternal signal. Panel B: when offspring favour mixtures that are more biased towards one phenotype (*z*_2_) relative to their mothers, the conflict is either won by offspring, mothers or neither of them, dependent on the relative strength of the costs of maladaptation in each environment. Panel C: when offspring favour mixtures that are less extreme relative to what is favored by their mothers, offspring win the conflict, as a fully informative maternal signal never results in more extreme mixtures than what is favored by offspring. Parameters: *d* = 0.1, *ℓ* = 0.5, *σ*_12_ = 0.1, *σ*_21_ = 0.25, *n* =1.

#### 3.2.1 Scenario 1: when offspring favour more extreme mixtures than their mothers, mothers typically win the conflict by providing partial information

When offspring favour more extreme mixtures of phenotypes relative to their mothers (as in Figure 1A), maternal signals often evolve to be partially informative to offspring, particularly when survival costs of a maladapted offspring (*c*_1_, *c*_2_) are large in both environments (Figure 2A, right corner). In this case, offspring are selected to rely on maternal information, as the alternative results in substantial costs due to local maladaptation. However, by limiting the amount of information about the local environment, mothers force offspring to increase their level of bet-hedging, thus resulting in a less extreme mixture of offspring phenotypes that coincides with the maternal optimum. (see Figure 3A). An example of such a partially informative signaling strategy is given in Figures 4A,B (see Figure S6 for a corresponding individual-based simulation).

**Figure 4.**
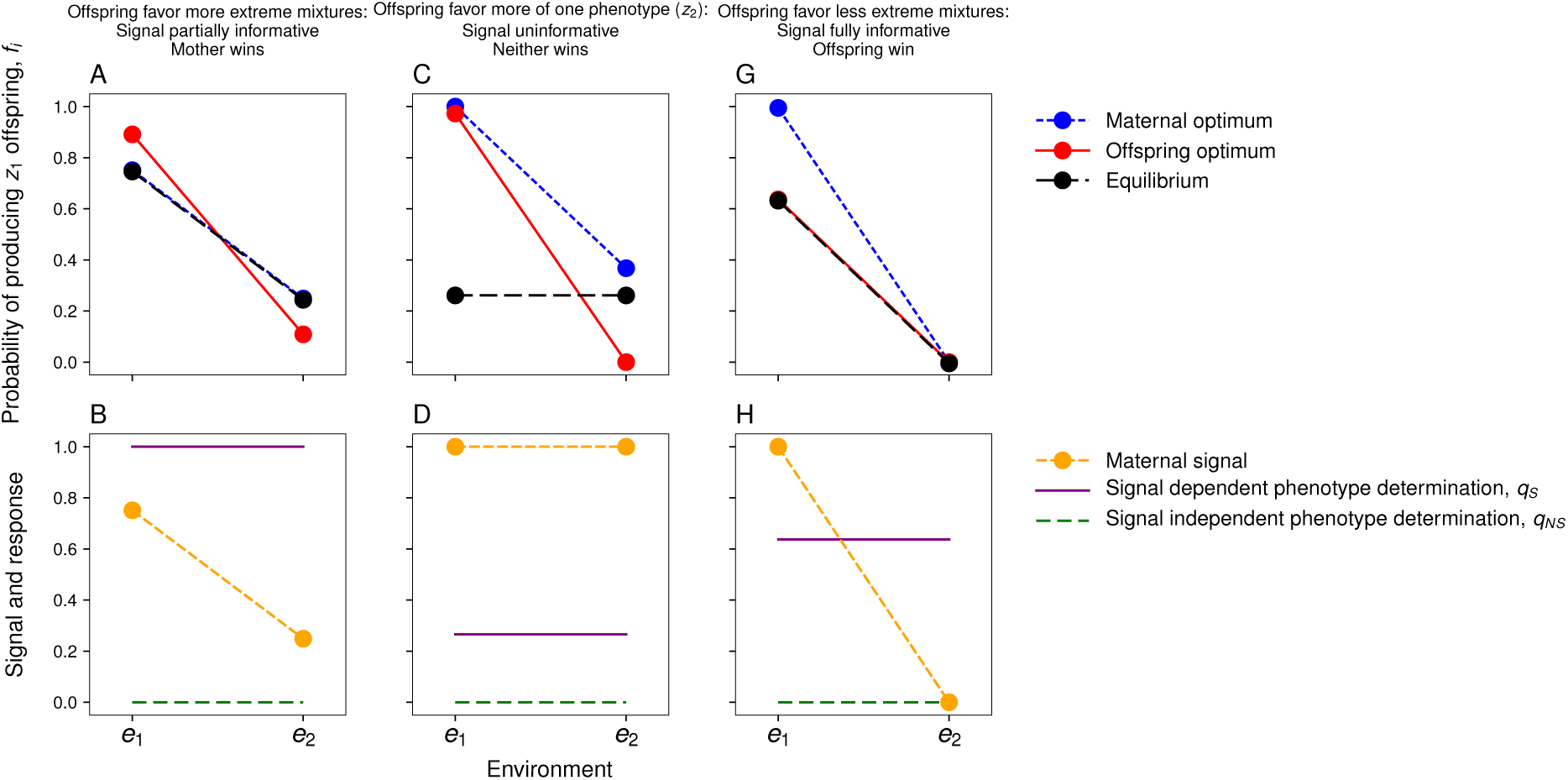
Examples of phenotype determination strategies (top row) and resulting signalling strategies (bottom row). Panels A, B: when offspring favour more extreme mixtures of phenotypes than mothers, mothers evolve only partially informative signals (panel B). As offspring only receive a limited amount of environmental information, offspring produce less extreme phenotypic mixtures, and mothers win the conflict (panel A). Panels C, D: when offspring favour phenotypic mixtures that are more biased towards one phenotype (*z*_2_), uninformative maternal signals commonly evolve (panel D), so that neither parents nor offspring win the conflict (panel C). Panels E, F: when offspring favour less extreme mixtures of phenotypes than mothers, mothers evolve fully informative signals (panel F). As a result, offspring obtain complete environmental information, resulting in offspring winning the conflict (panel C). Parameters: *d* = 0.1, *ℓ* = 0.5, *σ*_12_ = *σ*_21_ = 0.15, *n* = 1. Specific parameters for the different panels: A, B: *c*_1_ = *c*_2_ = 0.95, *β*_1_ = *β*_2_ = *γ*_1_ = *γ*_2_ = 1; C, D: *c*_1_ = 0.5, *c*_2_ = 0.8, *β*_1_ = *β*_2_ = 1, *γ*_1_ = *γ*_2_ = 2; E, F: *c*_1_ = 0.2, *c*_2_ = 0.8, *β*_1_ = *γ*_2_ = 1, *β*_2_ = *γ*_1_ = 2. The scenario in panels C, D where offspring favour phenotypic mixtures that are more biased towards one phenotype (*z*_2_) is further highlighted in Supplementary Figure S5.

Note, however, that coevolution between maternal signals and offspring responsiveness can also lead to alternative outcomes: when the cost of maladaptation is large in one environment, but small in the other, maternal signals evolve to become uninformative (white regions in Figure 2A), as mothers favour the exclusive production of a single offspring phenotype (the one having the highest costs of maladaptation) across the two environments. Conversely, when costs of maladaptation are modest and of similar magnitude in both environments, parental and offspring optima align, leading parents to evolve signals that are fully informative to offspring (black region in Figure 2A).

#### 3.2.2 Scenario 2: when offspring favour more of one phenotype, then typically neither side wins, and signalling often breaks down

As described above, when one phenotype is more costly to produce than the other, offspring favour mixtures in both environments that are more biased towards the more expensive phenotype (here *z*_2_) than do mothers (see the battleground in Figure 1B). By far the commonest outcome for this type of battleground is that signaling breaks down (light grey areas in Figure 2B), thus resulting in unconditional offspring phenotype determination strategies. Who wins the conflict now starts to depend on the relative costs of maladaptation (Figure 3B): when survival costs of maladaptation are high in environment *e*_1_, yet very low in environment *e*_2_ (white regions in Figure 3B), both parents and offspring favour the production of a single phenotype (*z*_1_; which matches the most severe environment) across both environments, so conflict is absent. When costs of maladaptation in environment *e*_2_ are slightly larger, however, offspring born in environment *e*_2_ favour the production of costly *z*_2_ offspring, while mothers still favour the production of *z*_1_ offspring in both environments. However, in the presence of an uninformative signal, offspring are forced to play an unconditional strategy which results in the exclusive production of *z*_1_ offspring in both environments, so that mothers can be said to win the conflict (light grey area in Figure 3B).

For even higher costs of maladaptation in environment *e*_2_ in Figure 3B, mothers again favour the exclusive production of *z*_1_ offspring across both environments (see Figure 4C, D for a detailed example). However, offspring now favour the production of a mixture of both *z*_1_ and *z*_2_ offspring in the absence of any maternal information, so that the resolution is one in which neither parent nor offspring wins the conflict (black region in Figure 3B). Finally, when costs of maladaptation are high in environment *e*_2_, but not in environment *e*_1_ (right part in Figure 2B), mothers too now start to favour the production of some costly *z*_2_ offspring in environment *e*_2_ (see Figure 4E, F for a detailed example). However, as offspring favour a much larger proportion of *z*_2_ offspring in environment *e*_2_ than mothers do, mothers only provide a partially informative signal to offspring (dark grey area in Figure 2B). The resulting uncertainty leads to less extreme proportion of *z*_2_ offspring in environment *e*_2_, but also leads to the production of some *z*_2_ offspring in environment *e*_1_. Consequently, again neither parents or offspring can be said to win the conflict (see Figure 4E).

#### 3.2.3 Result 3: when offspring favour less extreme mixtures than their mothers, offspring typically win the conflict, with mothers providing full information

When phenotype *z*_2_ is more costly to produce in environment *e*_1_, while phenotype *z*_1_ is more costly to produce in environment *e*_2_, mothers favour mixtures that are more extreme than offspring do (see the corresponding battleground in Figure 1C). Regarding the resolution of the conflict, Figure 2C shows that maternal signals either evolve to be fully informative, or that offspring evolve to be unresponsive to maternal signals (barring narrow regions in which signals are partially informative). In addition, Figure 3C shows that, for this configuration of maternal production costs, there is a substantial region where conflict is absent. However, when conflict occurs, offspring win the conflict as a result of these fully informative signals.

A more detailed example is shown in Figures 4G, H: to avoid the production of offspring that are more costly in terms of maternal resources, mothers favour extreme mixtures consisting only of *z*_1_ offspring in environment *e*_1_ and only of *z*_2_ offspring in environment *e*_2_ (blue dotted line in Figure 4C). Offspring, however, favour a less extreme mixture of phenotypes (red solid line). Mothers are then selected to provide offspring with the maximum amount of information, as this yields mixtures of phenotypes that are closest to what is favoured by the mother. By contrast, would mothers reduce the information content of the maternal signal, they would only select offspring to produce even less extreme mixtures that are even further away from the maternal optima. Hence, provided with complete environmental information, offspring can attain their respective optima in each environment (black dotted and red lines overlap in Figure 4C).

## 4 Discussion

While our model is general in formulation, it is applicable to many concrete types of maternal effect, whenever mothers can influence the cues available to offspring in a way that potentially reflects the local environment. A biological example of such a mechanism is the provision of different concentrations of a maternal hormone or small RNAs to young in different environments (Groothuis & Schwabl, 2008; Meylan *et al.*, 2012; Liebers *et al.*, 2014). In this kind of situation, mothers can provide offspring with an informative signal by varying hormone or small RNA concentrations markedly across environments, or by contrast, withold information by providing more similar concentrations of the same hormone across environments. The same reasoning applies to heritable epimutations (Heard & Martienssen, 2014), where strong vs weak differences in DNA methylation of gametes between environments reflect a strongly vs weakly informative maternal signal.

Our main conclusion is that parent-offspring conflict can have a significant impact on the evolution of informative maternal effects, even when offspring are unconstrained in their responses. The key feature of our model that leads to this outcome is that parents are allowed to adopt an imperfectly informative signalling strategy, and to ‘skew’ offspring responses towards their preferred outcome by independently adjusting the probabilities with which they give or withold signals in each environment. When mothers can potentially manipulate offspring in this way, we find that parent-offspring conflict often leads to a partial or even a complete breakdown in information transfer at equilibrium (just as it can do in models of signalling of need by offspring to their parents, Johnstone & Godfray, 2002). Consider, for instance, the case in which parents favour a higher proportion of a cheaper phenotype among their young, compared to that favoured by their offspring, and in which they do so regardless of the local environment. Under these conditions, it is hard for informative maternal signals to persist. If offspring take advantage of this information by responding to such a signal, an individual mother can always ‘push’ her young closer towards her own optimum by misrepresenting the state of the environment, and signalling in a way typical of local conditions that elicit a higher proportion of the cheaper phenotype. Consequently, we conclude that parent-offspring conflict may provide a powerful explanation for the apparent weakness of transgenerational plasticity in nature (for reviews see Uller *et al.*, 2013; Heard & Martienssen, 2014)

The possibility in our model for parents to independently adjust the probabilities with which they give or withhold signals in each environment explains the contrast between our results and those of Uller & Pen (2011). In their pioneering study of the impact of parent-offspring conflict on the evolution of maternal signals, Uller & Pen (2011) found that offspring typically evolve to be highly sensitive to maternal information about the state of the environment, regardless of any discrepancy between maternal and offspring optima. Their main model, however, assumes that offspring are provided with a highly discrete signal *m_i_* that is tied to a particular patch type *e_i_.* Consequently, even the slightest divergence between maternal signals in each environment (i.e., *m_i_* ≠ *m_j_*) provides offspring with perfect information about the environment. In other words, mothers are only able to withhold information to offspring when they are able to hold the signal *m_i_* exactly equal to *m_j_*, which would require considerable canalization in the face of mutation and drift in either signal.

In a supplementary model, Uller & Pen (2011) also briefly analyzed the evolution of maternal assessment errors, where making an error in environment *e_i_* implies that offspring are provided with signal *m_j_* rather than signal *m_i_.* However, they found that these maternal errors do not evolve. A key assumption of this extended model is, however, that the error is constrained to be identical across both environments: hence, a nonzero error only evolves when the advantage of providing offspring with a wrong signal in environment *e_i_* outweighs the disadvantage of providing offspring with a wrong signal in environment *e_j_* too. As a consequence, there is no scope for parents to independently adjust the probability of a signal being given in each environment, and so no possibility for parents to skew offspring responses in their own favour by misrepresenting the environment in a biased manner. Overall, this raises the question which mechanism is more realistic: are maternal effects indeed constrained as in the model of Uller & Pen (2011), or is there sufficient flexibility as required by the current model? Because maternal effects like hormones are often highly flexible (e.g., Müller *et al.*, 2004; Krist & Munclinger, 2015) and characterized by continuous (rather than discrete) variation across environments (e.g., Pavitt *et al.*, 2014; Lessells *et al.*, 2016), we suggest that the scope for maternal manipulation, as described by the current model, is likely to be substantial. Perhaps one way through which both the model of Uller & Pen (2011) and the current one can be reconciled, however, is when offspring are able to enforce honesty in maternal signals, so that maternal manipulation is not possible. It is therefore important that future models assess the evolutionary potential for enforcing honesty in maternal signals, whether this indeed leads to offspring winning the conflict, and how transgenerational plasticity is affected by honest signals. Yet, we emphasize that the potential for manipulation cannot be ruled out a priori for any signaling system (Dawkins & Krebs, 1978; Laidre & Johnstone, 2013), and maternal signals are no exception to this.

Another key conclusion of our model is that a reduction in maternal information transfer does not necessarily imply that either mothers (or offspring) win the conflict. Rather, the outcome of the conflict typically depends on the nature of the disagreement between mothers and young (see Figure 1), which depends on the specific trait that is studied. We suggest that scenarios in which mothers favour a more even mixture of phenotypes than do offspring (see Figure 1A) are more likely to result in partially informative signals and mothers winning the conflict (see Figure 3A). This type of outcome is particularly likely when alternative offspring phenotypes impose roughly similar costs on their mothers. One possible example is when individuals bet-hedge defences against multiple stressors, as they do when resistance to one strain of parasite trades off against resistance to another strain (strain-specific immunity: Little *et al.*, 2003; Schmid-Hempel, 2005. While resistance in such contexts is often studied in the context of heterozygosity (e.g., Penn *et al.*, 2002), an accumulating number of studies have shown that parasite resistance is, in part, influenced by transgenerational effects (Little *et al.*, 2003; Boulinier & Staszewski, 2008; Rechavi, 2014; Pigeault *et al.*, 2016). Our model predicts that parents would be selectively favored to produce more even mixtures of offspring resistant to one parasite strain versus another, while offspring themselves favour resistance against the parasite that is commonest in current local environment. In contexts like these, we would expect that mothers only provide their offspring with limited amounts of information about local parasite prevalence (leading to limited amounts of transgenerational plasticity - Uller, 2008; Holeski *et al.*, 2012), resulting in mothers winning the conflict.

For those traits for which offspring always favour overproduction of the costliest phenotype relative to mothers (see Figure 1B), it is more difficult to predict who wins the conflict: dependent on the parameters involved, either the offspring, the mother, or neither wins the conflict (Figure 3B). More important, however, is our finding that maternal information transfer can completely break down in this scenario, resulting in an absence of transgenerational plasticity (Figure 2B), which is particularly likely to occur when costs of maladaptation are modest. We believe that the battleground depicted in Figure 1B applies to numerous traits that have been previously studied in the context of parent-offspring conflict. For example, when the trait in question is offspring size (Smith & Fretwell, 1974), offspring will always favour a larger size than mothers themselves (Parker & Macnair, 1978; Einum & Fleming, 2000; Parker *et al.*, 2002; Kuijper & Johnstone, 2012). Similarly, when the trait in question is germination or diapause, offspring favour earlier germination than do their mothers because this enhances their probability of survival, while mothers favour later germination because this reduces competition with siblings (Ellner, 1986). Finally, in the context of sex allocation, mothers favour overproduction of the cheaper sex (Trivers, 1974; Kuijper & Pen, 2014), or the sex that is least affected by local competition (Werren *et al.*, 2002; Pen, 2006; Wild & West, 2009).

For those traits for which offspring favour less extreme mixtures relative to their mothers (see Figure 1C), we predict that it is nearly always offspring who win the conflict, unless costs of maladaptation are very high (Figure 3C). More importantly, we predict that maternal signals are often fully informative in such scenarios. However, we struggle to think of specific traits that are likely to fit these assumptions.

Summing up, we make two main, testable predictions. First, since parent-offspring conflict will often partially or completely destabilise maternal signalling, we predict that informative maternal effects are more likely to evolve, and to exert stronger effects, where conflict between parent and offspring is less pronounced. In other words, informative maternal effects should be strongest in contexts of female monogamy or when females reproduce asexually. Such a prediction could, for example be tested among closely related species with different mating systems, as is the case for the nematode genus *Caenorhabditis* (Fierst *et al.*, 2015; Teotónio *et al.*, 2017). Second, given that the impact of parent-offspring conflict depends upon the nature of the disagreement between parent and offspring, we predict that at least partially informative maternal effects are most likely to evolve or persist (even in the face of parent-offspring conflict) when different phenotypes impose similar costs to mothers (e.g., bet-hedging against different strains of parasites), we would expect partially informative signals to evolve. By contrast, breakdown of maternal signalling is a more likely outcome for traits in which one offspring phenotype is more costly to mothers than other offspring phenotypes (e.g., dispersal, sex allocation, germination), particularly when the costs of local maladaptation are modest.

## Acknowledgements

BK has been funded by an EPSRC 2020 Science fellowship (grant number EP/I017909/1) and a Leverhulme Trust Early Career Research Fellowship (ECF 2015-273). RAJ was funded by an EPSRC sandpit grant on transgenerational effects, grant number EP/H031928/1. This work has made use of the Carson computing cluster at the Environment and Sustainability Institute at the University of Exeter. In addition, the authors acknowledge the use of the UCL Legion High Performance Computing Facility (Legion@UCL) and associated support services in the completion of this work. The Dutch Academy of Arts and Sciences (KNAW) and the Lorentz Centre at the University of Leiden, the Netherlands funded a workshop on nongenetic effects that contributed to this article.

